# Differential Transcript Usage Analysis Incorporating Quantification Uncertainty Via Compositional Measurement Error Regression Modeling

**DOI:** 10.1101/2020.05.22.111450

**Authors:** Scott Van Buren, Naim Rashid

## Abstract

Differential transcript usage (DTU) occurs when the relative transcript abundance of a gene changes between different conditions. Existing approaches to analyze DTU often rely on computational procedures that can have speed and scalability issues as the number of samples increases. In this paper, we propose a new method, termed *CompDTU*, that utilizes compositional regression to model transcript-level relative abundance proportions that are of interest in DTU analyses. This procedure does not suffer from speed and scalability issues due to the relative computational simplicity, making it ideally suited for DTU analysis with large sample sizes. The method also allows for the testing of and controlling for multiple categorical or continuous covariates. Additionally, many existing approaches for DTU ignore quantification uncertainty present in RNA-Seq data, where prior work has shown that accounting for such uncertainty may improve testing performance. We extend our *CompDTU* method to incorporate quantification uncertainty using bootstrap replicates of abundance estimates from *Salmon* and term this method *CompDTUme*. Through several power analyses, we show that *CompDTU* improves sensitivity and reduces false positive results relative to existing methods. Additionally, *CompDTUme* results in further improvements in performance over *CompDTU* with sufficient sample size for genes with high levels of quantification uncertainty while maintaining favorable speed and scalability.

## 1. Introduction

Alternative splicing, the process by which a single gene may encode for multiple proteins, occurs naturally across cell types and species. It is a vital mechanism that allows cells to adapt to their environment by determining the specific set of coding regions in a gene that may be expressed (Kelemen et al., 2013). Differential transcript usage (DTU) is a special case of alternative splicing in which a gene’s relative transcript abundance (RTA) profile, the proportion of total gene-level expression that arises from each of the gene’s transcript isoforms, differs between conditions (Soneson et al., 2016). In contrast to typical differential gene or transcript isoform expression analyses, differences in the total gene expression levels are not of primary interest in DTU analyses. An example of DTU may be illustrated by the scenario where the total gene expression is the same between conditions A and B, but transcript isoform 1 is the predominantly expressed transcript in condition A while transcript isoform 2 is the predominantly expressed transcript in condition B. Specific RTA profiles have been identified in many diseases such as cancer (Scotti and Swanson, 2015), where the functional impact of DTU is only beginning to be understood (Climente-González et al., 2017).

Several methods have been recently proposed to detect the presence of DTU between conditions, such as *DRIMSeq* (Nowicka and Robinson, 2016). *DRIMSeq* estimates the RTA profile for each gene in each condition via a Dirichlet-multinomial model, given transcript-level read counts estimated by expression quantification using programs such as *Salmon* (Patro et al., 2017) or *kallisto* (Bray et al., 2016). A likelihoodratio test is performed to evaluate the null hypothesis that the mean RTA profiles pertaining to conditions *l* = 1,…, *L*, denoted **π**_*l*_, are equal across the *L* conditions, indicating no DTU. However, the complexity of the method can lead to speed and scalability issues as the number of samples increases, which may hinder the method’s practical applicability. Recent developments in sequencing techniques that have promised massive reductions in sequencing costs per sample (Alpern et al., 2019) enable greatly increased sample sizes and have made scalability of methods to large sample sizes an increasingly important characteristic.

In addition, transcript isoform-level expression abundances provided by common RNA-seq quantification programs are not directly observed and are instead computationally inferred (Patro et al., 2017; Bray et al., 2016). Specifically, RNA-seq reads may align to exons that are shared between multiple transcript isoforms, and therefore unique assignment of a single read to a single transcript isoform in such cases is difficult (Mortazavi et al., 2008). To address this difficulty, methods such as *Salmon* utilize an EM framework to estimate the conditional expectation of unobserved transcript abundances given a set of RNA-seq reads and known transcript isoforms pertaining to each gene. As a result, statistical uncertainty in these estimates, deemed “inferential variance” (Zhu et al., 2019), is present. This uncertainty may be further compounded by factors such as low overall gene expression or higher sequence similarity of transcript isoforms belonging to a given gene (Pimentel et al., 2017).

Prior work has shown that ignoring inferential variance when present may bias the estimation of the true biological variance of expression, affecting downstream statistical inference (Pimentel et al., 2017; Zhu et al., 2019). For example, *sleuth* (Pimentel et al., 2017) is a method for differential gene and transcript isoform expression analysis that decomposes total observed expression variation across samples into true biological variance and inferential variance. The latter term is estimated using “bootstrap replicates” or “inferential replicates”, which are abundance estimates that are computed from bootstrap samples from the estimated model in *kallisto*. Their results show that this decomposition improves the sensitivity and specificity of detecting differentially expressed genes and transcripts. *Salmon* can similarly generate bootstrap replicates by sampling counts for each equivalence class (with replacement) and rerunning the offline inference procedure (either the EM or VBEM algorithm) for each bootstrap sample (Patro et al., 2017). In general, inferential replicates can facilitate estimation of the within-sample transcript expression quantification uncertainty, which may improve the accuracy of between-sample biological variance.

*RATs* (Relative Abundance of Transcripts) is another method designed to identify DTU (Froussios et al., 2019) and utilizes inferential replicates from *kallisto* or *Salmon. RATs* applies G-tests of independence (McDonald, 2014) to test for DTU between conditions and repeats this testing procedure on each inferential replicate. A gene is classified as having DTU if the null hypothesis of no DTU is rejected for *x*% of inferential replicates across samples (where *x* is user-specified) and if the gene meets specific filtering criteria. This repeated testing across inferential replicates and samples may cause speed and scalability issues as the number of biological samples or inferential replicates increase. Most significantly, the choice of *x* is arbitrary. *RATs* additionally is unable to accommodate more than two conditions and is unable to control for or conduct significance tests with respect to predictors other than condition. *BANDITS* (Tiberi and Robinson, 2020) is another method designed to identify DTU that accounts for quantification uncertainty at the equivalence class level, treating the allocation of reads to specific transcripts as parameters that are sampled using Markov Chain Monte Carlo (MCMC) methods. While the method shows favorable performance in terms of sensitivity and specificity relative to existing methods, the method is demonstrated to have a gene-level computation time that is between six and 12 times slower than *DRIMSeq* (Tiberi and Robinson, 2020), sharing the speed and scalability issues present for *DRIMSeq*. Additionally, *BANDITS* is unable to control for or conduct significance tests with respect to predictors other than condition.

To mitigate the limitations of existing methods, we propose an efficient, scalable, and flexible compositional regression modeling framework (Pawlowsky-Glahn and Buccianti, 2011) to detect DTU based upon the application of the isometric log-ratio transformation (*ilr*) (Egozcue et al., 2003) to the set of observed sample-level RTA profiles for each gene. Our proposed compositional regression approach builds upon standard multivariate regression methods such as MANOVA, facilitating significant speed and scalability improvements relative to existing approaches while also maintaining better performance in detecting DTU. We additionally extend our compositional regression model to allow the incorporation of inferential replicates via a measurement error in the response modeling approach (Buonaccorsi, 2010), which shows improved performance in detecting DTU with similar computational efficiency. Our approach is additionally able to handle arbitrary design matrices, which may be helpful when adjusting for potential confounders, testing with respect to an arbitrary number of conditions, or evaluating the relevance of one or more continuous covariates of interest. We demonstrate the utility of our approach compared to existing methods through several extensive simulation studies.

## 2. Methods

### 2.1. Preprocessing

Define *T_ijg_* as the transcript expression measurement for sample *i, i* = 1,…, *N*, and transcript isoform *j, j* = 1,…, *D_g_*, pertaining to some gene *g* with *D_g_* known transcript isoforms, where *g* = 1,…, *G*. To simplify notation, we drop the subscript *g* pertaining to gene hereafter. Commonly used quantities for RNA-seq transcript isoform expression include Transcripts Per Million (TPM) (Wagner, Kin and Lynch, 2012), which provides a transcript-length and library size normalized estimate of expression for each transcript isoform within a sample. This correction facilitates comparison of abundance estimates across transcript isoforms within the same gene, as the estimated number of RNA-seq reads mapping to a gene or transcript isoform is often correlated with its length (Wagner, Kin and Lynch, 2012). Then, the RTA specific to transcript isoform *j* can be defined as 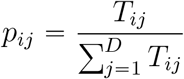 and the RTA profile for a given gene can be denoted as the vector ***p**_i_* = (*p*_*i*1_,…, *p_iD_*), where 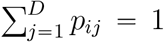, *p_ij_* ≥ 0 for all *i* and *j*. In other words, *p_ij_* pertains to the observed fraction of the total gene expression in sample *i* belonging to transcript isoform *j*. We also may wish to model ***p**_i_* with respect to some vector of subject-level predictors ***x**_i_*. In the DTU setting, ***x**_i_* may contain categorical variables corresponding to condition or treatment along with other arbitrary predictors of interest.

Given this setup, ***p**_i_* is inherently compositional in nature, and therefore lies on a standard *D*-part Aitchison simplex such that 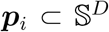, with 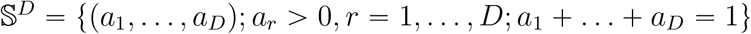 (Pawlowsky-Glahn and Buccianti, 2011). Direct regression modeling of compositional outcomes has been shown to present multiple difficulties (Aitchison, 1982), and therefore transformations are typically utilized to facilitate downstream statistical inference. Common transformations include the additive log-ratio transformation (*alr*) transformation (Aitchison, 1986), defined as *alr*(***p**_i_*) = (log{*p*_*i*1_/*p_iD_*},…, log{*p*_*i*(*D*−1)_/*p_iD_*}) where transcript isoform *D* is utilized as the reference isoform. However, the *alr* transformation is not an isometric transformation, where distances between different elements of ***p**_i_* pre-transformation are not preserved post-transformation, potentially leading to inaccurate significance testing results (Pawlowsky-Glahn and Buccianti, 2011). In addition, post-transformation values may differ depending on which transcript isoform is selected as the reference (Egozcue et al., 2003), where the choice of a particular transcript isoform as the reference is arbitrary in DTU analysis.

In contrast, the *clr* transformation (Aitchison, 1986) is defined as *clr*(***p**_i_*) = (log{*p*_*i*1_/*g*(***p**_i_*)},…, log{*p_iD_*/*g*(***p**_i_*)}), where *g*(***p**_i_*) = (*p*_*i*1_*p*_*i*2_ … *p_iD_*)^1/*D*^ is the geometric mean of ***p**_i_*. As a result, the *clr* avoids one of the drawbacks of the *alr* by utilizing *g*(***p**_i_*) as a common “reference.” The *clr* transformation is an isometric transformation, but *clr*-transformed vectors sum to zero, and therefore covariance matrices estimated from *clr*-transformed vectors are singular (Pawlowsky-Glahn and Buccianti, 2011). This result hinders the application of common multivariate testing approaches to *clr* transformed values (Rencher, 2002).

To avoid these limitations, Egozcue et al. (2003) proposed the use of an isometric logratio transformation (*ilr*), which has been previously used in analyses of microbiome data (Washburne et al., 2017; Silverman et al., 2018). This *ilr* transformation modifies the *clr* such that

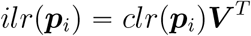

where ***V*** is a (*D* – 1) × *D* matrix whose rows form an orthonormal basis of 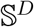. As its name suggests, the *ilr* transformation is an isometric transformation from the *D*-part Aitchison simplex 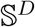 onto 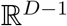 (Egozcue et al., 2003). This vector ***y**_i_* = *ilr*(***p**_i_*) = (*y*_*i*1_,…, *y*_*i*{*D*−1}_) is thus comprised of (*D* – 1) individual *ilr* coordinates and is commonly assumed to follow a (*D* – 1) dimensional multivariate normal distribution with full-rank covariance matrices (Pawlowsky-Glahn and Buccianti, 2011; van den Boogaart and Tolosana-Delgado, 2013). These properties facilitate the application of an efficient regression-based approach (Pawlowsky-Glahn and Buccianti, 2011) to DTU analysis, which we elaborate on in the next section. Note that the *ilr* transformation cannot accommodate any elements of ***p**_i_* being exactly equal to zero. We modify values of ***p**_i_* that are equal to or close to zero to facilitate downstream estimation and testing. Details are given in Section 1 of the Supplementary Material (Van Buren and Rashid, 2020).

### 2.2. Compositional Regression Model

We specify the following compositional regression model (*CompDTU*) utilizing the *ilr*-transformed coordinates to test for DTU. Letting *K* = *D* – 1, we assume ***y**_i_* = ***x**_i_**β*** + ***ϵ**_i_*, where ***y**_i_* is the vector of *ilr* coordinates (Section 2.1), ***x**_i_* is an 1 × *S* vector of covariates, ***β*** is the *S* × *K* matrix of regression coefficients, and ***ϵ**_i_* is the 1 × *K* vector of error terms. Additionally, define ***β**_k_* as the set of coefficients corresponding to the *k^th^ ilr* coordinate for *k* = 1,…, *K* and ***β*** = (***β***_1_,…, ***β**_k_*,…, ***β**_K_*). Note that ***x**_i_* is fixed with respect to *k*. We assume that 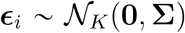 and therefore 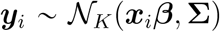, with 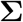 being the *K* × *K* covariance matrix. Then, given the above assumptions, the log-likelihood of this model is given by

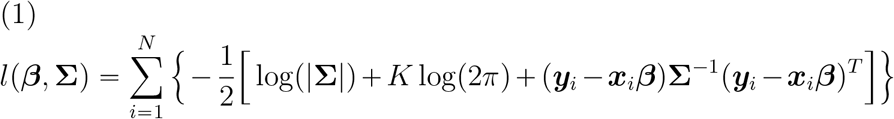

If we let 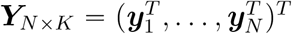 and 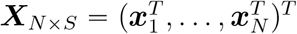, then the Maximum Likelihood Estimate (MLE) of ***β*** is given by 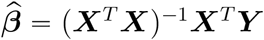 and the MLE of 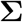 is given by 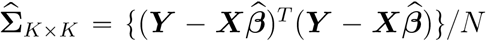 (Konishi, 2014). The predicted RTA for sample *i* is then given by 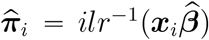, where *ilr*^-1^ denotes the inverse *ilr* transformation. If we assume that ***X*** pertains to the design matrix representing a categorical predictor (such as condition), this model is equivalent to the standard MANOVA model (Muller and Fetterman, 2003). As mentioned previously, we repeat this procedure for each gene.

#### 2.2.1. Hypothesis Testing

Hypothesis testing to evaluate the presence of DTU across conditions involves the evaluation of the null hypothesis *H*_0_: ***π***_1_ = … = ***π**_L_*, which tests for equivalence of the condition-specific mean RTA. This is equivalent to testing *H*_0_: ***β**_COND_* = **0**, where ***β_COND_*** is the subset of ***β*** corresponding to condition. For example, considering only two conditions, we can define the design matrix as ***X*** = (***x***_1_,…, ***x**_N_*)^*T*^, where ***x**_i_* = (1, 0) if sample *i* is in condition 1 and ***x**_i_* = (1,1) if sample i is in condition 2. Then, we can define *β*_*k*0_ as the intercept for *ilr* coordinate *k* (corresponding to the condition 1 mean) and *β*_*k*1_ as the difference in the means between condition 2 and condition 1 for *ilr* coordinate *k*. Then, ***β**_COND_* = (*β*_11_,…, *β*_11_,…, *β*_*K*1_), and ***β**_COND_* = 0 implies equality in the means across conditions such that ***π***_1_ = … = ***π**_L_*.

Several MANOVA test statistics are available to evaluate *H*_0_, such as the Pillai-Bartlett (Pillai, 1955; Bartlett, 1938), Wilks (Wilks, 1932), Hotelling-Lawley (Hotelling, 1951; Lawley, 1938) and Roy’s greatest root (Potthoff and Roy, 1964) statistics. While all four give identical results in some cases, such as when only two condition levels are being tested (Rencher, 2002), we use the Pillai-Bartlett statistic as it has been shown to be more robust to departures from the homogeneity of covariance assumption that is needed by a MANOVA model than alternative approaches (Hand and Taylor, 1987). This test is based on an eigenvalue decomposition of 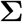 from (1), estimated separately under the null and alternative hypotheses.

Specifically, if we let 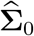 and 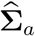 be the MLEs of 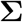 under *H*_0_ and *H_a_* respectively, we can define 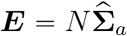 and 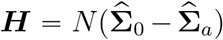. Then, the Pillai statistic *V* can be defined as

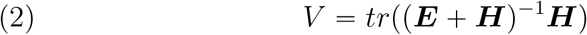

with *V* ~ *F*(*df*_1_, *df*_2_) under *H*_0_. To define *df*_1_ and *df*_2_, let *r* be the number of eigenvalues of **E**^−1^ **H**, *q* be the degrees of freedom of the hypothesis test, *t* = *min*(*r, q*), and *v* be the residual degrees of freedom corresponding to the alternative hypothesis. Then, *df*_1_ = *t*(*abs*(*r*–*q*)+*t*) and *df*_2_ = *t*(*v* – *r* + *t*). Note that the *df*_1_ and *df*_2_ specified here give an approximate *F* distribution for *V* under *H*_0_ that is valid for general settings, and exact values of *df*_1_ and *df*_2_ are available in some cases (Rencher, 2002).

This approach is applicable to arbitrary ***X***, which is helpful when the inclusion of additional categorical or continuous predictors into the model is desired. Also, 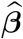 and 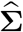 can be estimated from (1) under the null and alternative hypotheses in closed-form through efficient matrix-based operations (Section 2.2), and therefore this framework is extremely fast and scalable.

Rejecting the null hypothesis for the significance test of condition suggests the overall presence of DTU for a given gene. However, it is important to note that there is no direct one–to–one mapping between elements of ***p**_i_* and ***y**_i_* (van den Boogaart and Tolosana-Delgado, 2013), as the procedure described provides overall tests for association. To determine which pairs of transcript isoforms exhibit DTU following a significant overall test, one may restrict the comparison to pairs of transcript isoforms from the same gene and repeat the analysis procedure detailed in Sections 2.1 and 2.2. Given the sequential nature of this test and the number of pairwise tests performed, correcting for the multiple tests performed in sequence is necessary and can be done using the *stageR* procedure (Van den Berge et al., 2017).

### 2.3. Measurement Error Compositional Regression Model

To account for quantification uncertainty, we now propose a modification of our proposed *CompDTU* model. This new approach, which we call *CompDTUme*, utilizes inferential replicates of TPM values instead of the standard TPM point estimates utilized by *CompDTU*. Let *N* be the total number of samples, *M* be total number of inferential replicates sampled, and *K* = *D* – 1 again be the total number of *ilr* coordinates for an arbitrary gene with *D* transcript isoforms. Letting *i* = 1,…, *N* index sample, *j* = 1,…, *M* index the inferential replicate number, and *k* = 1,…, *K* index the *ilr* coordinate number, we define ***y**_ij_* = (*y*_*ij*1_,…, *y_ijK_*) to be the set of *ilr*-transformed coordinates pertaining the *j*’th inferential replicate for sample *i*. We would like to incorporate all *M* inferential replicates for all *N* samples into the model, and to do this we can define the overall *NM* × *K* response matrix ***Y**_ME_* as 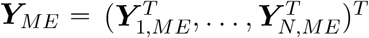, where 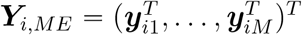. Additionally, let ***X**_ME_* be the *NM* × *S* design matrix formed by repeating each row of ***X*** from Section 2.2 *M* times such that ***X**_ME_* = (***x***_1,*ME*_,…, ***x**_N,ME_*)^*T*^, where ***x**_i,ME_* is the *M* × *S* matrix of repeated covariates for sample *i*. Denote 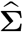 as the MLE of the total covariance matrix from (1) replacing ***Y*** with ***Y**_ME_* and ***X*** with ***X**_ME_*.

We assume that the total covariance for sample *i* can be decomposed into the sum of between-sample and within-sample components (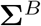 and 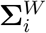 respectively) such that the total covariance for sample *i* is 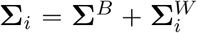, similar to methods used in longitudinal data analysis (Fitzmaurice, Laird and Ware, 2004). The estimate of the between-subjects covariance matrix 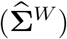 can be obtained using a multivariate extension of the moment-based estimator for variance under measurement error with normally distributed response values presented in Buonaccorsi (2010) such that

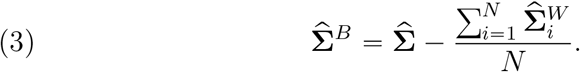

Here, 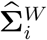 is estimate for the *K* × *K* within-sample covariance matrix for sample *i*, denoted as 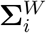. We may obtain element (*a, b*) of its estimate 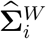 by calculating

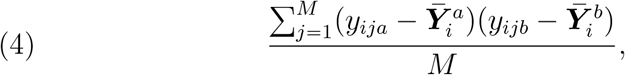

where 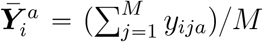 and 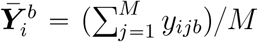 utilize the information from the *M* inferential replicates from sample *i*.

We propose conducting hypothesis tests using the Pillai-Bartlett test statistic as discussed in Section 2.2.1, now utilizing 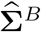 instead of 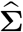. We will show that this modification of the Pillai-Bartlett procedure will provide improved testing performance by removing the effect of inferential covariance from the total covariance matrix. We evaluate this claim in simulations presented in Section 4.

Two types of inferential replicates are available from *Salmon* to use with *CompDTUme*, bootstrap replicates and Gibbs replicates, described previously in Section 1. It is unclear which approach to generating inferential replicates is preferred in general. However, we found significant autocorrelation between Gibbs inferential replicates in certain transcripts under default settings (see Section 2 of the Supplementary Material (Van Buren and Rashid, 2020)). Such autocorrelation is commonly observed in samples obtained using MCMC procedures. For these reasons, we use bootstrap replicates for *CompDTUme* as they are independent by design.

## 3. Example Dataset

To compare computation times of *DRIM-seq, RATs, BANDITS, CompDTU*, and *CompDTUme* to detect DTU across conditions, we utilize the E-GEUV-1 data from the Geuvadis consortium (Lappalainen et al., 2013). This dataset contains quality controlled RNA-Seq data from lymphoblastoid cell lines pertaining to 462 unrelated individuals from five different human populations in the 1000 Genomes Project (Abecasis et al., 2012). Samples were quantified using *Salmon* (Patro et al., 2017), where 100 bootstrap inferential replicates were generated per sample, and the *tximport* package was utilized to import the *Salmon* results into *R* (Soneson, Love and Robinson, 2016). Pre-filtering of genes and transcript isoforms has been shown to greatly improve performance in DTU applications (Soneson et al., 2016). We utilize gene and transcript level filters from *DRIMSeq* using recommendations from Love, Soneson and Patro (2018) to facilitate comparisons across different methods. Each method was run on the 7,522 genes that passed filtering (with a total of 24,624 transcript isoforms passing filtering across those genes), and we use population membership as the condition variable in this analysis. Thus, results test for differences in RTA across the five populations such that *H*_0_: ***π***_1_ = … = ***π***_5_, where ***π**_l_* is the population-level mean RTA for a given gene for population *l* = 1,…, 5. Details of the computational and filtering options used are given in Section 3 of the Supplementary Material (Van Buren and Rashid, 2020).

Table 1 shows the computation time comparisons for the E-GEUV-1 dataset. We run the comparisons on the full 462 sample dataset across all five populations as well as on a subset of 20 samples that contains 10 randomly chosen samples from the “CEU” and “GBR” populations, respectively. We define our design matrix as described in Section 2 using reference cell coding for condition and follow the described testing framework.

**Table 1.**
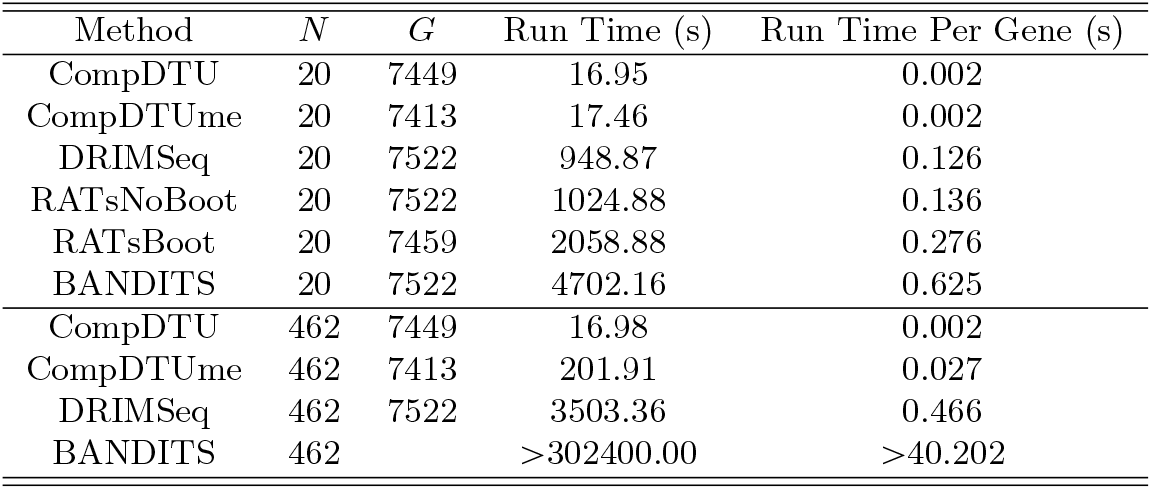
Computation time comparisons based on the E-GEUV-1 data across five conditions (N = 462) and a reduced subset spanning two conditions. G corresponds to the number of genes with non-missing significance results for each method from the 7,522 genes that passed filtering. Computation times are given in seconds.

| Method | $N$ | $G$ | Run Time (s) | Run Time Per Gene (s) |
| --- | --- | --- | --- | --- |
| CompDTU | 20 | 7449 | 16.95 | 0.002 |
| CompDTUme | 20 | 7413 | 17.46 | 0.002 |
| DRIMSeq | 20 | 7522 | 948.87 | 0.126 |
| RATsNoBoot | 20 | 7522 | 1024.88 | 0.136 |
| RATsBoot | 20 | 7459 | 2058.88 | 0.276 |
| BANDITS | 20 | 7522 | 4702.16 | 0.625 |
| CompDTU | 462 | 7449 | 16.98 | 0.002 |
| CompDTUme | 462 | 7413 | 201.91 | 0.027 |
| DRIMSeq | 462 | 7522 | 3503.36 | 0.466 |
| BANDITS | 462 |  | >302400.00 | >40.202 |

Our results show that the *CompDTU* and *CompDTUme* methods have significantly shorter computation times per gene than the other methods. In addition, we find that computation time per gene for the *CompDTU* method does not noticeably increase when applied to the full *N =* 462 sample E-GEUV-1 dataset relative to the *N =* 20 sample subset due to its computational efficiency. We do observe an increase in computation time for the *CompDTUme* method relative to *CompDTU* but the total time is still significantly less than *DRIMSeq* and additionally incorporates bootstrap replicates. *BANDITS* run on the full 462 sample dataset did not complete after 84 hours, and therefore we set its run equal to 84 hours and assume *G* = 7, 522 in Table 1. We were unable to evaluate *RATs* on the full 462 sample dataset because the method cannot accommodate more than two conditions. Overall, both *CompDTU* and *CompDTUme* run significantly faster than the existing methods and both scale more efficiently with an increasing number of samples than other methods. See Section 3 of the Supplementary Material (Van Buren and Rashid, 2020) for details of computational options used for *DRIMSeq, RATs*, and *BANDITS*.

Because it is not known *a priori* which genes should truly show DTU across populations, computation of common metrics that benchmark the relative performance of each method is difficult. Instead, we utilize two simulation-based approaches discussed in the following section to compare the relative performance of each method in terms of their sensitivity and specificity to detect DTU, as well as to examine aspects such as type I error rates.

## 4. Simulation Studies

### 4.1. Simulated Multivariate Normal Outcomes

We directly compare the performance of the proposed *CompDTU* and *CompDTUme* methods across various conditions by simulating data directly from the multivariate normal distribution. First, denote 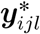 as the 1 × *K* vector of simulated *ilr* coordinates for condition *l* = 1,…, *L* for sample *i* = 1,…, *N*, and inferential replicate *j* = 1,…, *M* for a gene with *K ilr* coordinates. The number of sampled observations is the thus same in each condition and equal to *N*. We simulate data from the multi-variate normal distribution using mean and covariance estimated from gene ENSG00000002822.15 from the full 462 sample E-GEUV-1 data results discussed in Section 3. Specifically, let 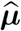 be the vector of mean *ilr*-transformed coordinates across all 462 samples, obtained by fitting an intercept-only *CompDTU* model to the data from the gene. Additionally, let 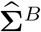 be the corresponding *K* × *K* estimated between-sample covariance matrix obtained by fitting an intercept-only *CompDTUme* model to the data pertaining to this gene.

To simulate 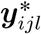, we first obtain a within-sample covariance matrix 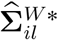 by sampling with replacement from the set 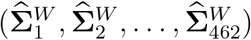, which pertains to each sample from the E-GEUV-1 data for gene ENSG00000002822.15. Then, assuming *L* = 2, 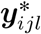 is sampled from 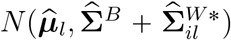, where 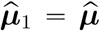 and 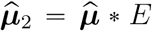. Here, *E* controls the effect size, inducing mean differences between the two conditions. Setting *E* = 1 enables the evaluation of type I error under no DTU. To decrease or increase the amount of measurement error in the simulation relative to the amount present in the observed data for the gene, the diagonal elements of 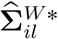 are multiplied by a multiplicative factor 0 < *H* < 1 or *H* > 1, respectively. The selected gene, ENSG00000002822.15, has four unique transcript isoforms that pass filtering (*D* = 4), and was chosen because it is representative of genes with moderately large count values. Therefore, for our simulation we assume *L* = 2 conditions, *K* = 3, *N* = 10, 26, 100, *M* = 50, 100, *H* = 0.01,1, 2, and *E* = 1,1.1,1.25,1.375,1.5,1.75, and 2.

Simulation results are shown in Table 2. Results pertaining to column *E* = 1 correspond to data generated under the null hypothesis of no DTU between conditions, and therefore pertain to the estimated type I error of each method. Our results show that *CompDTU* and *CompDTUme* both approximately preserve type I error. Slight inflation of type I error can be observed for *CompDTUme*, however this is mitigated by increasing the number of bootstrap replicates utilized in the model. In addition, accounting for quantification uncertainty via *CompDTUme* can result in large improvements in power over *CompDTU* for moderate effect sizes, especially as the level of measurement error increases. See Section 4 and Tables S2 and S3 in the Supplementary Material for these results as well as results from additional simulation scenarios (Van Buren and Rashid, 2020). An effect size of 1.50 in this context corresponds to a 4.25% change in the average RTA proportion of the major transcript isoform of the gene between two conditions. This 4.25% change corresponds to the 87th percentile of the same quantity observed across genes in the unmodified E-GEUV-1 data, making it a moderately large effect size based on this data.

**Table 2.**
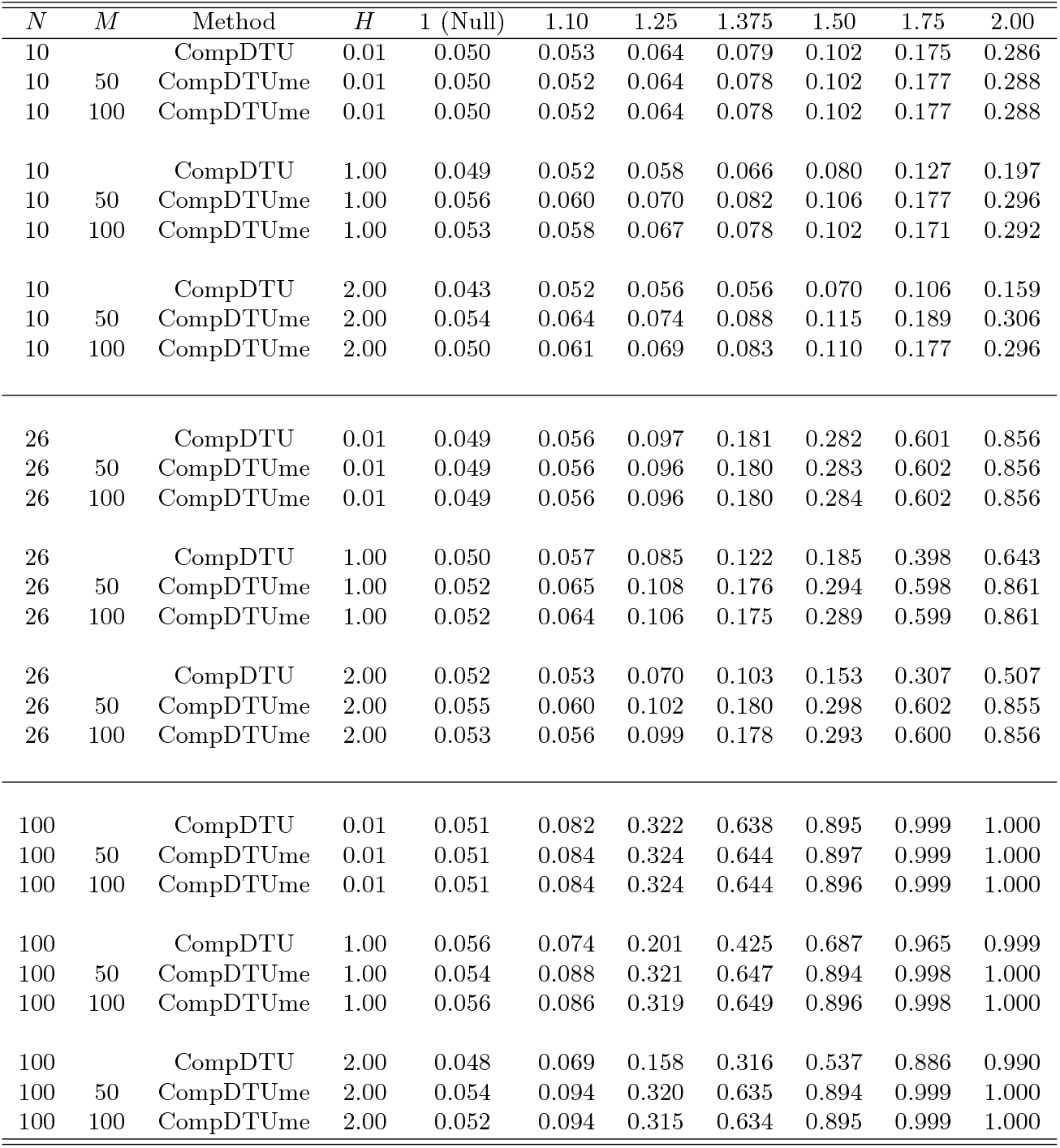
Power for CompDTU and CompDTUme across various simulation conditions. N pertains to the sample size per condition, and M pertains to the number of inferential replicates per sample. H reflects the relative amount of measurement error. The effect size E is given by the numerical column names and ranges from 1 to 2. All simulations assume L = 2 conditions and D = 4 transcript isoforms.

### 4.2. Permutation-Based Simulation from Real Data

#### 4.2.1. Procedure

To facilitate the comparison of our proposed methods to existing approaches, we perform a simulation study utilizing the E-GEUV-1 data described in Section 3. To perform this simulation, we randomly choose 50 samples from both the “CEU” and “GBR” populations for a total of 100 samples, and use these populations as the two conditions in our simulation. To induce a difference between the two conditions, we first randomly select half of the study genes (comprised of the 7,522 genes that passed filtering for the analysis described in Section 3) to show DTU across conditions. We then randomly generate 100 permutations of samples with respect to the “CEU” and “GBR” conditions. For each permutation, we induce a difference in the counts of the major transcript isoform between conditions among genes selected to show DTU. Specifically, for samples assigned to the “CEU” label, we multiply the counts of the major transcript isoform by a specific “change value” for genes randomly selected to show DTU. We then recompute the TPM values using the updated counts and sample-specific estimated transcript isoform lengths from *Salmon*. Modification is done on the counts instead of on the TPM values for ease of use with *DRIM-Seq* and *RATs*, both of which do not use TPM values. A change value of 1 corresponds to the null hypothesis of no DTU, and allows us to evaluate the type I error of each method across conditions. Change values greater than 1 induce a difference between the two conditions and allow us to evaluate sensitivity and specificity of each method under a known effect size (change value).

We apply the same permutation assignments and subsequent count modification procedure to each of the 100 sets of bootstrap replicates obtained from *Salmon* for each sample and gene. We varied the magnitude of the change value to evaluate the relative power of each method to detect DTU across different effect sizes, but only present results corresponding to a change value of 2 for brevity. This procedure is repeated on each of the 100 sample permutations across conditions, and results are pooled across genes and condition arrangements. We do not compare to *BANDITS* because of the lack of computational scalability and the method’s use of equivalence classes instead of count or TPM point estimates. See Section 5 of the Supplementary Material for additional information (Van Buren and Rashid, 2020).

#### 4.2.2. Results

Figure 1 shows histograms of *p*-values under the null hypothesis from the E-GEUV-1 permutation-based simulation. Histograms were generated after pooling results across all genes and condition permutations. This figure demonstrates that *p*-values from *CompDTU* and *CompDTUme* most closely follow a uniform distribution under the null hypothesis relative to other methods. A slight increase in type I error is observed for *CompDTU* and *CompDTUme*, but this increase is less than the corresponding increase observed for *DRIMSeq*, with the type I errors for the three methods equal to 0.056, 0.058, and 0.083, respectively. The results for *CompDTU* and *CompDTUme* are additionally very similar to the type I errors under the null hypothesis found in the simulation analysis presented in Table 2 with 100 samples and *H* = 1. In addition, we find that *RATs* filters out almost all genes, resulting in *p*-values for these genes being set to 1. This is in line with what is done in this scenario in Love, Soneson and Patro (2018), suggesting the default filtering procedure from *RATs* is overly aggressive.

**Fig 1:**
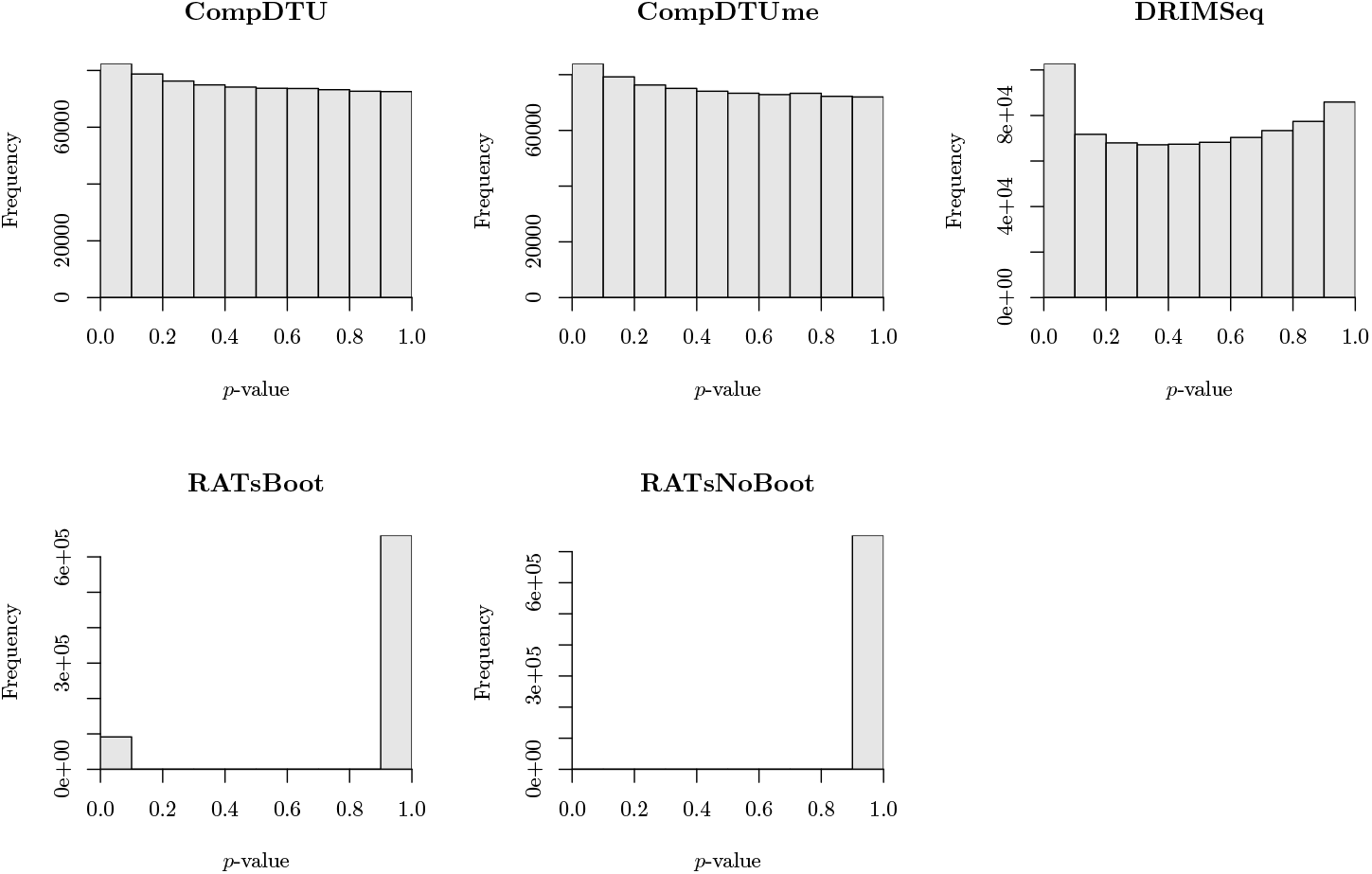
Histograms of *p*-values determined under the null hypothesis of no DTU, pooling results from all genes and permutations. The *RATs* methods by default utilize *p*-values from the G tests of independence as well as filtering based on effect size and reproducibility (Froussios et al., 2019). For *RATsBoot*, this reproducibility analysis incorporates bootstrap replicates, while for *RATsNoBoot* it does not. Following Love, Soneson and Patro (2018), we set *p*-values for genes that are filtered out by RATs to 1, which results in nearly all *p*-values having a value of 1.

Figure 2A plots the *p*-value significance threshold vs. the false pos-itive rate (FPR) under the null hypothesis, which is the proportion of genes that do not truly show DTU that incorrectly reject *H*0 to indicate significant DTU. In this figure, we again observe that *DRIMSeq* has higher type I error than *CompDTU* or *CompDTUme*. In contrast, *RATs* run with and without bootstrap replicates (*RATsNoBoot* and *RATsBoot* respectively) has inflated and deflated type I errors respectively. Figure 2B plots the estimated false discovery rate (eFDR) (Benjamini and Hochberg, 1995), which is the significance threshold used for *p*-values adjusted using the FDR method (Benjamini and Hochberg, 1995) against the True FDR, defined as the proportion of all significant tests that correspond to genes that do not truly show DTU at the given FDR threshold. This figure illustrates that *DRIMSeq* has slightly inflated True FDR values for eFDR cutoff values less than 0.05. This observation is not true for genes with transcripts that have low sequence overlap (Supplementary Figure S2A) (Van Buren and Rashid, 2020) and is worsened for genes with transcripts that have high sequence overlap (Supplementary Figure S2B). Sequence overlap in this context is defined as the proportion of exon base pairs that are shared between multiple transcripts within a specific gene. This quantity ranges between 0 and 1, where 0 indicates no exon base pairs are shared across different transcripts within a gene and 1 indicates all exon base pairs are shared. Estimation of transcript-level counts and TPMs is most difficult for genes with high levels of transcript isoform sequence similarity. *RATsBoot* and *RATsNoBoot* have inflated and deflated True FDR values across all genes respectively, with the former having an extremely inflated value of 0.464.

**Fig 2:**
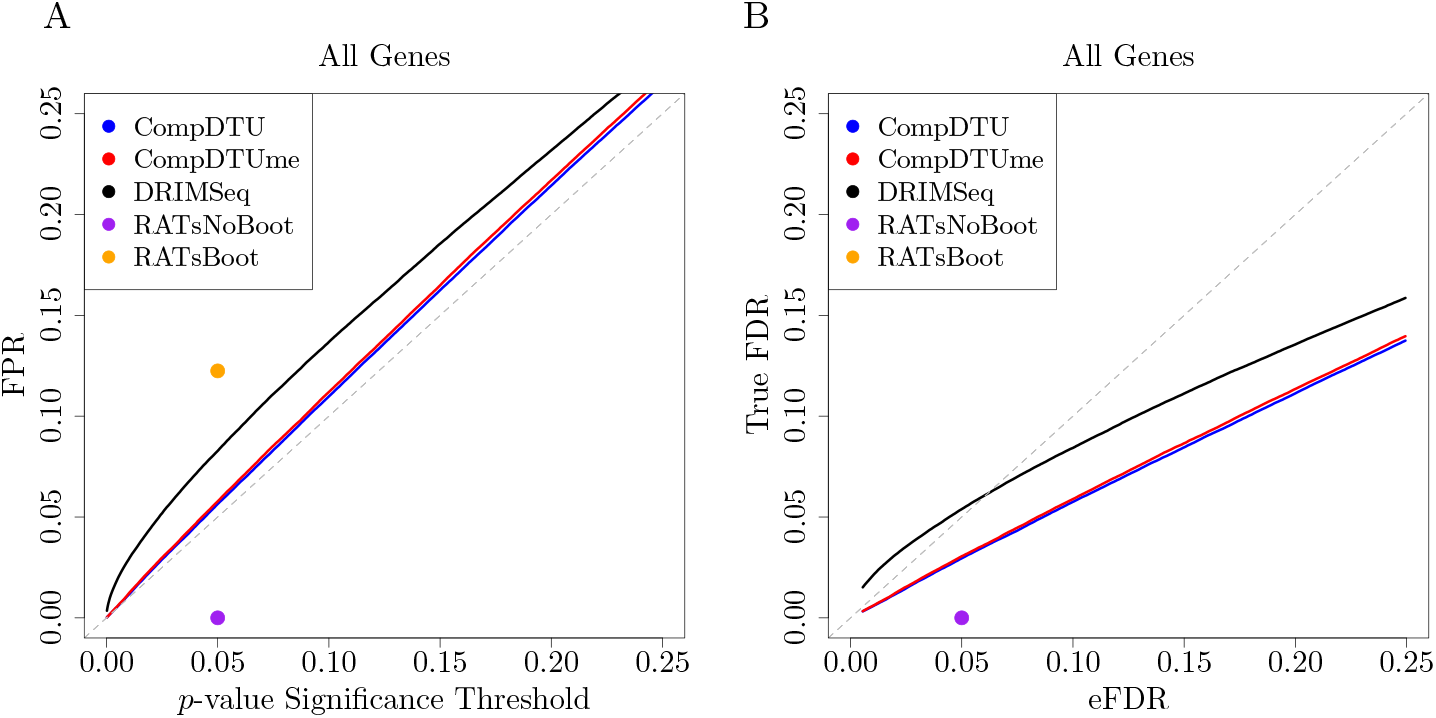
Comparisons of false positive/false discovery rates under the null hypothesis of no DTU (change value = 1, Panel A) and under a moderate effect size (change value = 2, Panel B) for the 7,522 genes that passed filtering. Panel A plots the unadjusted *p*-value threshold vs the false positive rate (FPR) under the null hypothesis for each method. Panel B plots the True FDR level for a given eFDR value for each method, where eFDR is the significance threshold chosen for *p*-values adjusted using the FDR. For *RATsBoot*, this reproducibility analysis incorporates bootstrap replicates, while for *RATsNoBoot* it does not. Following Love, Soneson and Patro (2018), we set *p*-values for genes that are filtered out by *RATs* to 1, which results in nearly all *p*-values having a value of 1. *RATsBoot* does not appear in Panel B because the True FDR observed is 0.464 at an eFDR value of 0.05. Note that for *RATs* only single points are able to be shown due to internal filtering procedures.

Figure 3 shows ROC curves for detecting DTU from data generated under a change value of 2. The results collectively show improved sensitivity by *CompDTU* and *CompDTUme* relative to *DRIMSeq*, and that *CompDTU* and *CompDTUme* have lower FPRs for a given eFDR cut-off than *DRIMSeq*. *RATsBoot* and *RATsNoBoot* show greatly reduced sensitivity compared to the other methods at an eFDR level of 0.05. This is likely due to the default filtering procedure to determine if a result is “reproducible” in *RATs* being too aggressive, resulting in only the most significant results remaining as significant. Love, Soneson and Patro (2018) reaches a similar conclusion, finding that the *RATs* procedure including bootstrap replicates from *Salmon* gives nearly identical results when using nominal FDR thresholds of 1%, 5%, and 10%, indicating only the most highly significant genes generally pass filtering.

**Fig 3:**
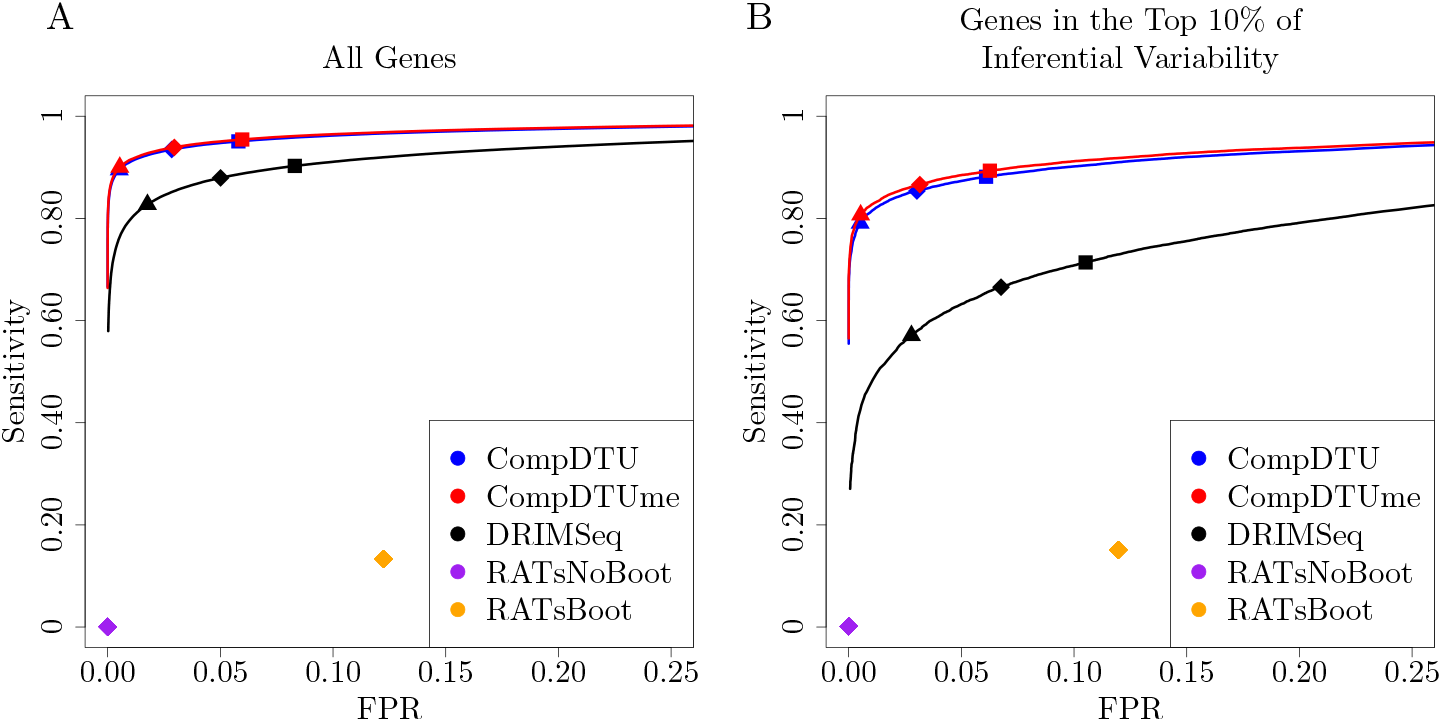
ROC Curves for detecting DTU across all 7,522 genes that pass filtering (Panel A) and the 752 genes in the top decile of gene-level inferential variability (Panel B). For details of how inferential variability is calculated, see Section 6 of the Supplementary Material (Van Buren and Rashid, 2020). A triangle, diamond, or square indicates the point at which the estimated FDR (eFDR) is 0.01, 0.05, or 0.10, respectively, where the eFDR is the FDR-adjusted *p*-value significance threshold.

We also find that *CompDTUme* shows similar performance to *CompDTU* across all genes (Figure 3A) but improved performance for the 752 genes that are in the top decile of genes in terms of inferential variability (Figure 3B). For details of how inferential variability is calculated, see Section 6 of the Supplementary Material (Van Buren and Rashid, 2020), but briefly we utilize the *InfRV* measure proposed in Zhu et al. (2019) to bin genes in terms of their inferential variability. This result corresponds with the simulation-based power analyses from Section 4.1 that demonstrate improved performance from *CompDTUme* relative to *CompDTU* as inferential variability increases.

Figure 4 shows boxplots of the RTA for the major transcript (ENST00000357955.6) of gene ENSG00000114738.10. This gene is in the top decile of inferential variability and was incorrectly not determined to show DTU by *CompDTU* (FDR adjusted *p*-value 0.061) and correctly determined to show DTU by *CompDTUme* (FDR adjusted *p*-value 0.006). Results are plotted across all 50 samples for each condition. The two boxplots on the left include a single estimate per sample that arises from the standard point estimates, and the two boxplots on the right include 100 estimates per sample that arise from bootstrap replicates. Results show that the median values for each condition are similar between the bootstrap and non-bootstrap values. However, use of the bootstrap replicates reveals a larger spread in the boxplot values for the “CEU” condition than is present when using the regular point estimates. This larger spread is reflective of the underlying quantification uncertainty present for the gene, and incorporation of this quantification uncertainty via *CompDTUme* results in *CompDTUme* reaching the correct decision of rejection while *CompDTU* does not. These results demonstrate that incorporation of bootstrap replicates via *CompDTUme* can improve performance over *CompDTU* when the level of inferential variability is high, agreeing with the results shown in Figure 3B and the simulation results presented in Table 2 and Tables S2 and S3 in the Supplementary Material (Van Buren and Rashid, 2020).

**Fig 4:**
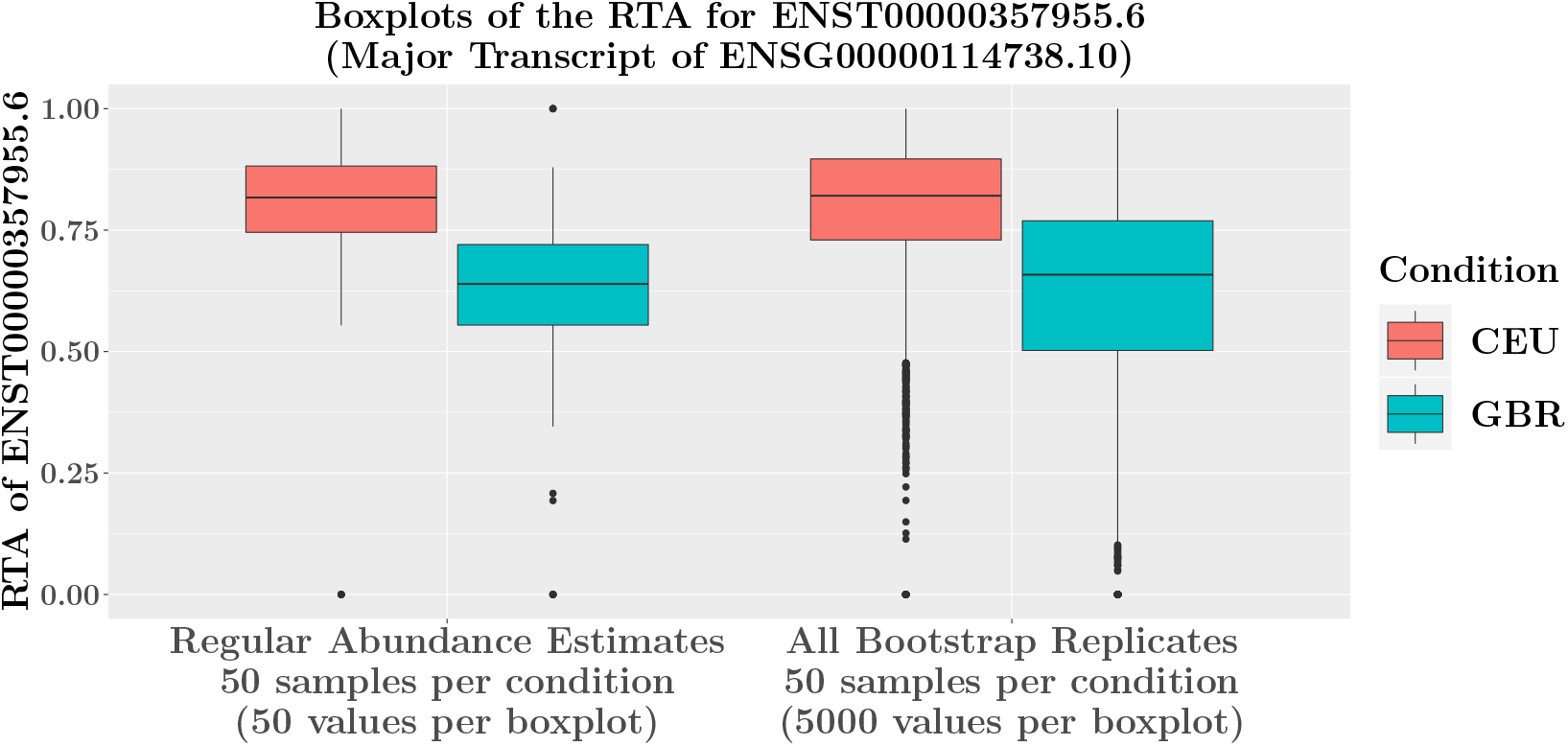
Boxplots of the RTA of the major transcript (ENST00000357955.6) of gene ENSG00000114738.10, which was incorrectly not determined to show DTU by *CompDTU* (FDR adjusted *p*-value 0.061) but correctly determined to show DTU (FDR adjusted *p*-value 0.006) by *CompDTUme*. Boxplots for the “Regular Abundance Estimates” are generated using the regular point estimates for all 50 samples for each condition, and boxplots for “All Bootstrap Replicates” are generated using data from all 100 bootstrap replicates for all 50 samples for each condition.

Results analogous to those presented in Figures 2 and 3 for a twenty-sample random subset of subjects (10 in each condition) are presented in Figures S3 and S4 respectively in Section 7 of the Supplementary Material (Van Buren and Rashid, 2020). Figure S3 demonstrates that *CompDTU* and *CompDTUme* successfully conserve FPR and FDR in a smaller sample size setting, while *DRIMSeq* has slightly inflated FPR and FDR. In addition to the expected decrease in sensitivity for each method resulting from each condition having only 10 samples, Figure S4 demonstrates improved performance of *CompDTU* and *CompDTUme* relative to *DRIMSeq* and *RATs*. Figure S4 also demonstrates that the inclusion of bootstrap replicates via *CompDTUme* results in slightly lower sensitivity than *CompDTU* with twenty samples for this simulation procedure, possibly due to inaccuracies in the estimate of the inferential covariance matrix given the small number of samples used. Additionally, Figure S5 plots ROC curves for results including *CompDTU* run using the mean of the bootstrap replicates instead of the usual point estimates (*CompDTUAbMeanBoot*) for both the 20 sample and 100 sample analyses. Results show that *CompDTUAbMeanBoot* performs similarly to *CompDTU* and *CompDTUme*. This result demonstrates that prior results showing superior performance from *CompDTUme* relative to *CompDTU* for genes with high levels of inferential variability are not driven by inherent differences between the point estimates and bootstrap samples.

Overall, these results, as well as those from Table 2 and Tables S2 and S3 from the Supplementary Material (Van Buren and Rashid, 2020) demonstrate bootstrap replicates can greatly improve performance when the number of biological samples is large enough to obtain an accurate estimate of the average inferential covariance. We recommend the use of *CompDTUme* if the total sample size is at least 25 and *CompDTU* otherwise.

## 5. Discussion

In this paper, we compare various methods to analyze DTU at the gene level, and propose a new approach, *CompDTU*, that is based on methods originally designed to analyze compositional data. We additionally extend our *CompDTU* method to incorporate information from inferential replicates from *Salmon* or *kallisto* to attempt to reduce the impact of quantification uncertainty on the final results and term this method *CompDTUme*. *CompDTUme* utilizes the covariance of the inferential replicates for a given gene averaged across samples and subtracts this covariance from the standard covariance estimate to obtain an improved estimate of the between-sample covariance matrix. This updated between-sample covariance matrix is then used with the Pillai test statistic to conduct hypothesis testing for DTU.

Table 1 shows that our proposed *CompDTU* and *CompDTUme* methods result in significant reductions in computation time compared to *RATs, DRIMSeq*, and *BANDITS*, and both methods scale with an increasing sample size much more efficiently than *DRIMSeq* and *BANDITS*. Additionally, our permutation-based power analysis from Section 4.2.2 finds that both *CompDTU* and *CompDTUme* result in increases in overall sensitivity and specificity compared to *RATs* and *DRIMSeq* and reduction in the FPR compared to *DRIMSeq*.

Our *CompDTUme* additionally shows increased sensitivity and specificity over the *CompDTU* method in our permutation-based power analysis from Section 4.2.2 for genes that are in the top 10% of inferential variability. Other datasets may have higher levels of measurement error than the E-GEUV-1 data, and simulation results presented in Table 2 and Tables S2 and S3 in the Supplementary Material (Van Buren and Rashid, 2020) show *CompDTUme* can greatly outperform *CompDTU* in the presence of increased inferential variability with a sufficient sample size. The ability of *CompDTUme* to accommodate bootstrap replicates in a computationally efficient manner can enable its use to protect against high levels of inferential variability that may be present in other datasets. Overall, given the results we have presented, we recommend the use of our *CompDTUme* method for DTU analysis when the number of samples is at least 25 (to ensure accurate estimates of the average inferential covariance needed to use *CompDTUme*) and *CompDTU* if not to maximize sensitivity to detect DTU while greatly reducing computation time relative to existing methods.

R code to run the proposed *CompDTU* and *CompDTUme* methods is available via an R package at https://github.com/skvanburen/CompDTUReg. This package includes all functions and sample code.

## SUPPLEMENTARY MATERIAL

### Online Supplement: Supplementary Material

(doi: COMPLETED BY THE TYPESETTER; .pdf). In Van Buren and Rashid (2020), we provide various additional details, including information about computational options used. We also provide additional simulation results.

## Notes

### Competing Interest Statement

The authors have declared no competing interest.

