## Supplementary Material for "Differential Transcript Usage Analysis Incorporating Quantification Uncertainty Via Compositional Measurement Error Regression Modeling"

May 17, 2020

#### 1 Correction of Lowly Expressed Transcripts in the CompDTU and CompDTUme Methods

The presence of zeros is a key problem in compositional data analysis, and many approaches have proposed to handle this issue (van den Boogaart and Tolosana-Delgado, 2013; Martín-Fernández et al., 2003). The nature of the *ilr* transformation implies that transformed coordinates are not defined for proportions that are exactly zero, and that small changes in proportions that are close to zero can lead to large changes in the values of their transformed coordinates (Egozcue et al., 2003). Proportions being zero or close to zero arises frequently in DTU analysis, as TPM values for a particular transcript isoform in a specific sample are frequently zero or close to zero, indicating no or biological low expression of that transcript isoform.

We overcome these issues by replacing any TPM value that is less than five percent of the total gene-level expression for the sample by five percent of this expression. Following the notation of the main text, let $T_{ij}$ be the TPM value for transcript isoform $j = 1, \ldots, D$ for sample $i = 1, \ldots, n$ within a given gene with $D$ transcript isoforms. If for transcript isoform $j$ we observe that $T_{ij} < 0.05 \times \sum_{j=1}^{D} T_{ij}$, we set $T_{ij} = 0.05 \times \sum_{j=1}^{D} T_{ij}$. The proportions and *ilr* coordinates are then recalculated according to the procedure detailed in Section 2.1 of the main text. This procedure results in relative transcript abundances (RTAs) being zero only when the total gene expression of the gene is equal to zero for the specific sample. In this case, the *ilr* coordinates will all be equal to zero, corresponding to the null hypothesis of no significant DTU across conditions. This approach is motivated by non-parametric zero-replacement methods described in Pawlowsky-

**Supplementary Table S1:** Average Autocorrelation Values at Lag 1 for Various Inferential Replicate Methods Across All Genes by Overlap Tertile for the SEQC Data

| Type | Thinning | LowerThird | MidThird | UpperThird |
|---|---|---|---|---|
| Gibbs | 16 | 0.048 | 0.089 | 0.145 |
| Gibbs | 100 | 0.011 | 0.023 | 0.047 |
| Boot | NA | -0.009 | -0.010 | -0.010 |

Glahn and Buccianti (2011). This type of procedure is simple, and similar procedures have been previously concluded to be a coherent and natural choice for the substitution of zeros and small non-zero values (Martín-Fernández and Thió-Henestrosa, 2006).

#### 2 Gibbs vs Bootstrap Inferential Replicates

Here we evaluate the use of both Gibbs and bootstrap samples from *Salmon* as inferential replicates. These evaluations were done using data from from the Sequencing Quality Control Project (SEQC) from two reference RNA samples, “A” and “B” (SEQC/MAQC-III Consortium, 2014). Sample A contains five technical replicates of the Strategene Universal Human Reference RNA (UHRR) and Sample B contains five technical replicates of the Ambion Human Brain Reference RNA (HBRR). We found that for certain transcripts the Gibbs inferential replicates showed high autocorrelation with the default thinning parameter value of 16, indicating that Gibbs replicates were saved every 16 Gibbs MCMC iterations (Figure S1). Thinning is often utilized in MCMC approaches to reduce correlation between draws from an MCMC chain to obtain approximately uncorrelated samples, where taking samples from MCMC iterations that are widely apart should decrease the correlation in save draws.

High levels of autocorrelation persisted for some transcripts even when increasing the thinning parameter to 100 (Table S1), and was especially the case for genes that have transcript isoforms that have high sequence similarity within a specific gene. For example, Table S1 shows the average autocorrelation values across all genes for replicate SRR950080 at a lag value of 1 for Gibbs samples using thinning values of 16 and 100 and for bootstrap samples, stratified by sequence overlap tertile. Sequence overlap in this context is defined as the proportion of exon base pairs that are in common

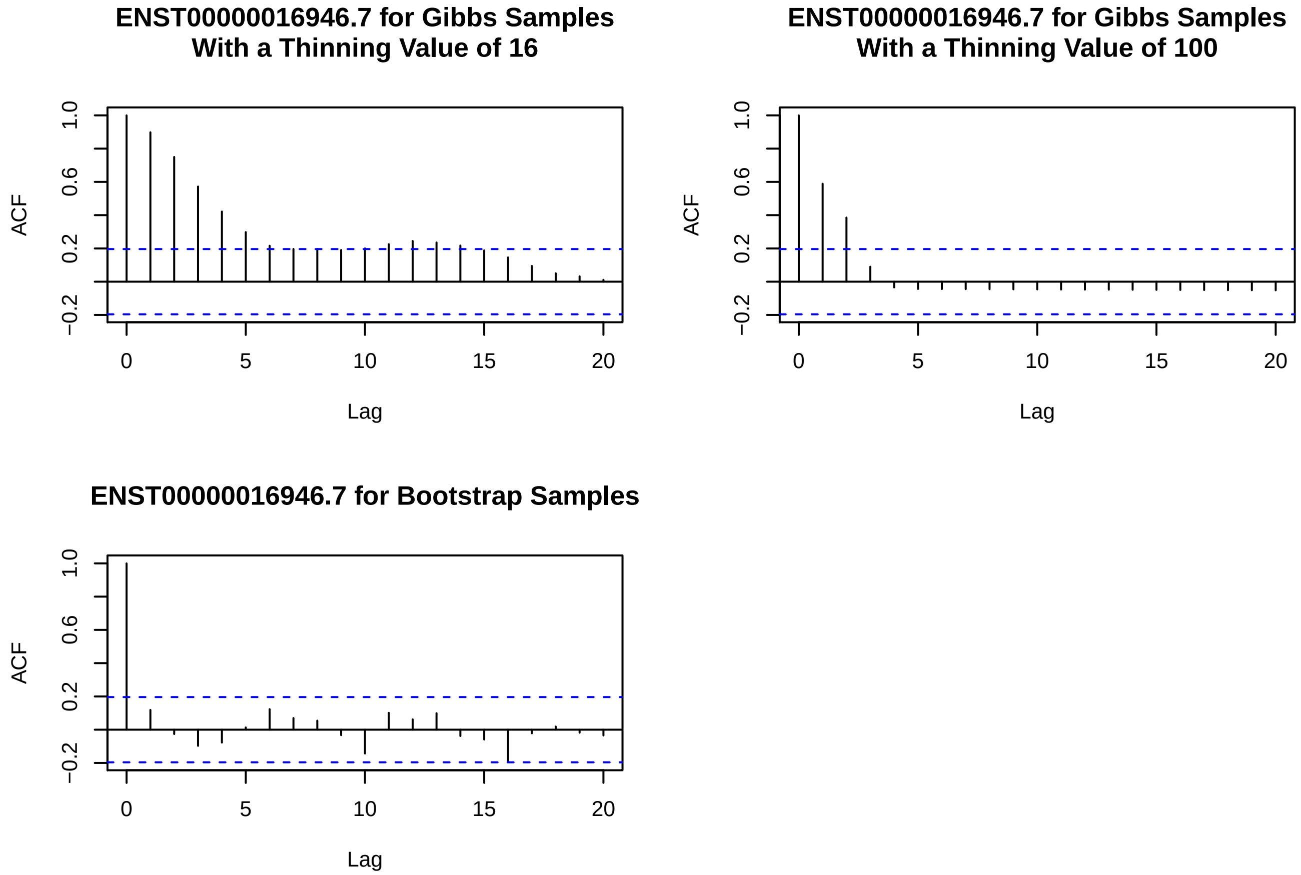

**Supplementary Figure S1:** Autocorrelation values for various lag values for transcript ENST00000016946.7 for Gibbs inferential replicates using thinning values of 16 and 100 and for bootstrap samples.

between more than 1 transcript isoform from a specific gene. It is thus calculated as a quantity between 0 and 1, with 0 indicating no exon base pairs are shared between transcript sequences for different transcripts within the gene and 1 indicating all exon base pairs are shared.

We find that the average autocorrelation at a lag value of 1 (i.e. for adjacent replicates) increases with the gene-level overlap tertile for a thinning value of 16 from 0.048 to 0.089 to 0.145. Increasing the thinning value to 100 reduces the average autocorrelation for each overlap tertile. In contrast, bootstrap samples show low average autocorrelation values at every overlap tertile. Figure S1 plots the autocorrelation of the inferential replicates for the transcript ENST00000016946.7, and shows Gibbs replicates with thinning values of 16 or 100 have much higher autocorrelation than those observed in bootstrap samples. Given these potential autocorrelation issues, we use bootstrap samples as inferential replicates instead of Gibbs samples.

### 3 Details of Various Computational Options

#### 3.1 *Salmon* Options

We performed all quantification steps in our paper using Salmon version 0.11.3 (Patro et al., 2017), with the index created using the GENCODEv27 annotation (Harrow et al., 2012; Frankish et al., 2018). The options used in the `salmon quant` step are: `--seqBias --gcBias --dumpEq --numBootstraps 100`, where the `--dumpEq` statement is utilized to save equivalence class information for use with the *BANDITS* method (Tiberi and Robinson, 2020).

#### 3.2 *tximport* Options

Following *Salmon* quantification, the data were loaded into `R` using version 1.14.0 of the `tximport` package (Soneson et al., 2016). We importantly generate estimated counts by scaling the TPM measurements up to the library size using the option `countsFromAbundance = "scaledTPM"`, as is recommended for DTU analyses in Love et al. (2018). All other options were left at their default values.

#### 3.3 *DRIMSeq* Filtering Parameters and Options

Our filtering method involves using filters built into *DRIMSeq* (Nowicka and Robinson, 2016) due to the fact that these are flexible and their use facilitates easy comparison to the method. First, let $n$ be the total sample size and $n.small$ be the number of samples in the condition with the fewest samples. As is done in Love et al. (2018), for a transcript to be kept in the dataset it has to have a count of at least 10 in at least $n.small$ samples, have a relative abundance proportion of at least 0.10 in at least $n.small$ samples, and the total count of the corresponding gene must be at least 10 in all $n$ samples.

*DRIMSeq* results were run using version 1.14.0 of the package (Nowicka and Robinson, 2016). We importantly use the option `add_uniform = TRUE` to allow estimates to be computed for genes with two features having all zeros for one group or genes with the last feature having all zeros for one group. However, we additionally noticed performance issues that were corrected by adding a count of 1 to all count values and thus added 1 to each count to stabilize results before running the *DRIMSeq* model. We additionally only fit the gene-level Dirichlet-multinomial models within the `dmFit` statement, and thus set `bb_model = FALSE`. All other options were left at their default values. Computation time results only include time required to run the `dmPrecision`, `dmFit`, and `dmTest` statements.

##### 3.4 *RATs* Parameters and Options

*RATs* was run using version 0.6.4 of the package (Froussios et al., 2019). We considered *RATs* both using and not using bootstrap samples from *Salmon* by setting `boot_data_A` and `boot_data_B` to be non-NULL in the former case and `count_data_A` and `count_data_B` to be non-NULL in the latter. All other options were left at their default values. Computation time results only include time required to run the `call_DTU` statement.

##### 3.5 *BANDITS* Parameters and Options

*BANDITS* was run using version 1.3.2 of the package (Tiberi and Robinson, 2020). We run the `prior_precision` statement to calculate an informative prior for the precision and use this with the `test_DTU` statement, as is highly recommended in the package's vignette. All other options were left at their default values. Computation time results only include the time required to run the `prior_precision` and `test_DTU` statements.

#### 4 Additional Simulation-Based Power Analysis Results

We evaluated the performance of our proposed approaches using the simulation-based procedure discussed in Section 4.1 of the main text under additional scenarios. Tables S2 and S3 show simulation-based power results for the *CompDTU* and *CompDTUme* methods when the amount of measurement error is multiplied by multiplicative factors of 0, 0.01, 0.50, 1.00, 1.50, 2, or 4 across varying sample sizes. This is done by multiplying the diagonal elements of the within-subject covariance matrix by the specified factor. An $H$ value of 0 corresponds to no measurement error, meaning *CompDTU* and *CompDTUme* should have identical power, which we observe.

Comparisons between *CompDTU* and *CompDTUme* reveal that *CompDTUme* can result in significant relative improvements in power relative to *CompDTU* while approximately conserving Type I error. Improvements in power from *CompDTUme* relative to *CompDTU* increase as $H$ increases, and the type I error observed for *CompDTUme* decreases as the number of replicates increases. For example, with the measurement error increased by a factor of 4 (such that $H = 4$) and 100 biological samples, results from Table S3 show that the use of 100 inferential replicates can increase the power with an effect size of 1.50 from 0.364 using *CompDTU* to 0.898 using *CompDTUme*.

Table S4 shows simulation-based power results for the *CompDTU* and *CompDTUme* methods when the between-subject variance terms are increased by a multiplicative factor of 1.50, 2, or 4. This table shows that Type I error values for both methods are not significantly affected by increased between-subject variance.

### 5 Additional Details About the Permutation-Based Power Simulation Procedure

As discussed in the main text, we set the change value for this power analysis to be 2. A change value of 2 corresponds to roughly the 90th percentile of the multiplicative difference in major transcript counts when comparing data averaged across samples from the "CEU" population to data averaged across samples for the "GBR" population in the unmodified E-GEUV-1 data, making it a moderately large effect size based on on the unmodified data. Note that the change values discussed here are modifications of the major transcript isoform abundance on the count scale, while the effect sizes used in the multivarite normal simulation-based power analyses are modifications of the *ilr* transformed RTAs derived from TPMs. Thus, the change values and effect sizes from the two different power simulations are not directly comparable.

### 6 Details about Gene-Level Inferential Variability Calculations

To estimate the gene-level variability, we utilize the InfRV measure proposed in (Zhu et al., 2019). This quantity is defined separately for each gene (or transcript) for each sample, and is given by:

$$InfRV = \frac{\max(s^2 - \mu, 0)}{\mu + 5} + 0.01$$

where $s^2$ and $\mu$ are the sample variance and mean values of the bootstrap (or Gibbs) samples for the given gene respectively. For our analysis, we calculate the $InfRV$ based on transcript-level values, and take the maximum of the transcript specific values for the gene as the final value for the gene for the current sample. The final overall value for the gene is taken as the average of the sample-specific values for the gene. This quantity is roughly independent of the range of the counts, and the quantities 5 and 0.01 are respectively added to stabilize the result and to ensure the final quantity is strictly positive. Note that $InfRV$ values are not used directly by *CompDTU* or *CompDTUme* but instead are used to generate the list of genes in the top 10% of inferential variability in Figure S4 and Figure 3B from the main text.

### 7 Additional Permutation-Based Power Simulation Results

#### 7.1 ROC Curve Results for 20 Sample Permutation-Based Analysis

Figures S3 and S4 plot results corresponding to Figures 2 and 3 in the main text for an analysis run on 20 total samples (10 randomly chosen samples across each of the "CEU" and "GBR" conditions) with 100 bootstrap replicates per sample. These results are discussed in Section 4.2.2 in the main text.

#### 7.2 *CompDTU* ROC Curve Results Run On The Mean of Bootstrap Samples

Figure S5 plots ROC curves for results including *CompDTU* run using the mean of the bootstrap samples instead of the usual point estimates (*CompDTUAbMeanBoot*). Results show that *CompDTUAbMeanBoot* performs similarly to *CompDTU* and *CompDTUme* across all genes.

**Supplementary Table S2:** Power for the *Comp* and *CompDTUme* Methods with Increased Measurement Error Variance. The effect size $E$ is given by the numerical column names and ranges from 1 to 2.

| $n$ | $M$ | Method | $H$ | 1 (Null) | 1.10 | 1.25 | 1.375 | 1.50 | 1.75 | 2.00 |
|---|---|---|---|---|---|---|---|---|---|---|
| 10 | | CompDTU | 0.00 | 0.047 | 0.047 | 0.067 | 0.078 | 0.101 | 0.177 | 0.291 |
| 10 | 50 | CompDTUme | 0.00 | 0.047 | 0.047 | 0.067 | 0.078 | 0.101 | 0.177 | 0.291 |
| 10 | 100 | CompDTUme | 0.00 | 0.047 | 0.047 | 0.067 | 0.078 | 0.101 | 0.177 | 0.291 |
| 10 | | CompDTU | 0.01 | 0.050 | 0.053 | 0.064 | 0.079 | 0.102 | 0.175 | 0.286 |
| 10 | 50 | CompDTUme | 0.01 | 0.050 | 0.052 | 0.064 | 0.078 | 0.102 | 0.177 | 0.288 |
| 10 | 100 | CompDTUme | 0.01 | 0.050 | 0.052 | 0.064 | 0.078 | 0.102 | 0.177 | 0.288 |
| 10 | | CompDTU | 0.50 | 0.049 | 0.052 | 0.060 | 0.074 | 0.094 | 0.146 | 0.226 |
| 10 | 50 | CompDTUme | 0.50 | 0.053 | 0.055 | 0.063 | 0.083 | 0.105 | 0.181 | 0.295 |
| 10 | 100 | CompDTUme | 0.50 | 0.053 | 0.054 | 0.062 | 0.081 | 0.104 | 0.178 | 0.292 |
| 10 | | CompDTU | 1.00 | 0.049 | 0.052 | 0.058 | 0.066 | 0.080 | 0.127 | 0.197 |
| 10 | 50 | CompDTUme | 1.00 | 0.056 | 0.060 | 0.070 | 0.082 | 0.106 | 0.177 | 0.296 |
| 10 | 100 | CompDTUme | 1.00 | 0.053 | 0.058 | 0.067 | 0.078 | 0.102 | 0.171 | 0.292 |
| 10 | | CompDTU | 2.00 | 0.043 | 0.052 | 0.056 | 0.056 | 0.070 | 0.106 | 0.159 |
| 10 | 50 | CompDTUme | 2.00 | 0.054 | 0.064 | 0.074 | 0.088 | 0.115 | 0.189 | 0.306 |
| 10 | 100 | CompDTUme | 2.00 | 0.050 | 0.061 | 0.069 | 0.083 | 0.110 | 0.177 | 0.296 |
| 10 | | CompDTU | 4.00 | 0.046 | 0.049 | 0.052 | 0.057 | 0.065 | 0.086 | 0.122 |
| 10 | 50 | CompDTUme | 4.00 | 0.068 | 0.073 | 0.083 | 0.100 | 0.130 | 0.206 | 0.323 |
| 10 | 100 | CompDTUme | 4.00 | 0.062 | 0.064 | 0.075 | 0.092 | 0.119 | 0.193 | 0.306 |
| 26 | | CompDTU | 0.00 | 0.049 | 0.062 | 0.098 | 0.177 | 0.294 | 0.597 | 0.861 |
| 26 | 50 | CompDTUme | 0.00 | 0.049 | 0.062 | 0.098 | 0.177 | 0.294 | 0.597 | 0.861 |
| 26 | 100 | CompDTUme | 0.00 | 0.049 | 0.062 | 0.098 | 0.177 | 0.294 | 0.597 | 0.861 |
| 26 | | CompDTU | 0.01 | 0.049 | 0.056 | 0.097 | 0.181 | 0.282 | 0.601 | 0.856 |
| 26 | 50 | CompDTUme | 0.01 | 0.049 | 0.056 | 0.096 | 0.180 | 0.283 | 0.602 | 0.856 |
| 26 | 100 | CompDTUme | 0.01 | 0.049 | 0.056 | 0.096 | 0.180 | 0.284 | 0.602 | 0.856 |
| 26 | | CompDTU | 0.50 | 0.050 | 0.056 | 0.084 | 0.145 | 0.227 | 0.481 | 0.737 |
| 26 | 50 | CompDTUme | 0.50 | 0.052 | 0.062 | 0.102 | 0.176 | 0.294 | 0.592 | 0.853 |
| 26 | 100 | CompDTUme | 0.50 | 0.052 | 0.061 | 0.100 | 0.174 | 0.295 | 0.591 | 0.853 |
| 26 | | CompDTU | 1.00 | 0.050 | 0.057 | 0.085 | 0.122 | 0.185 | 0.398 | 0.643 |
| 26 | 50 | CompDTUme | 1.00 | 0.052 | 0.065 | 0.108 | 0.176 | 0.294 | 0.598 | 0.861 |
| 26 | 100 | CompDTUme | 1.00 | 0.052 | 0.064 | 0.106 | 0.175 | 0.289 | 0.599 | 0.861 |
| 26 | | CompDTU | 2.00 | 0.052 | 0.053 | 0.070 | 0.103 | 0.153 | 0.307 | 0.507 |
| 26 | 50 | CompDTUme | 2.00 | 0.055 | 0.060 | 0.102 | 0.180 | 0.298 | 0.602 | 0.855 |
| 26 | 100 | CompDTUme | 2.00 | 0.053 | 0.056 | 0.099 | 0.178 | 0.293 | 0.600 | 0.856 |
| 26 | | CompDTU | 4.00 | 0.045 | 0.049 | 0.062 | 0.086 | 0.112 | 0.211 | 0.345 |
| 26 | 50 | CompDTUme | 4.00 | 0.055 | 0.062 | 0.113 | 0.190 | 0.308 | 0.595 | 0.860 |
| 26 | 100 | CompDTUme | 4.00 | 0.052 | 0.060 | 0.106 | 0.182 | 0.304 | 0.592 | 0.862 |

**Supplementary Table S3:** Power for the *Comp* and *CompDTUme* Methods with Increased Measurement Error Variance. The effect size $E$ is given by the numerical column names and ranges from 1 to 2.

| $n$ | $M$ | Method | $H$ | 1 (Null) | 1.10 | 1.25 | 1.375 | 1.50 | 1.75 | 2.00 |
|---|---|---|---|---|---|---|---|---|---|---|
| 50 | | CompDTU | 0.00 | 0.051 | 0.068 | 0.166 | 0.336 | 0.566 | 0.921 | 0.997 |
| 50 | 50 | CompDTUme | 0.00 | 0.051 | 0.068 | 0.166 | 0.336 | 0.566 | 0.921 | 0.997 |
| 50 | 100 | CompDTUme | 0.00 | 0.051 | 0.068 | 0.166 | 0.336 | 0.566 | 0.921 | 0.997 |
| 50 | | CompDTU | 0.01 | 0.047 | 0.068 | 0.170 | 0.345 | 0.564 | 0.912 | 0.995 |
| 50 | 50 | CompDTUme | 0.01 | 0.048 | 0.067 | 0.171 | 0.347 | 0.565 | 0.914 | 0.995 |
| 50 | 100 | CompDTUme | 0.01 | 0.047 | 0.067 | 0.171 | 0.348 | 0.566 | 0.914 | 0.995 |
| 50 | | CompDTU | 0.50 | 0.050 | 0.067 | 0.131 | 0.270 | 0.445 | 0.826 | 0.975 |
| 50 | 50 | CompDTUme | 0.50 | 0.052 | 0.070 | 0.164 | 0.343 | 0.571 | 0.920 | 0.995 |
| 50 | 100 | CompDTUme | 0.50 | 0.052 | 0.071 | 0.164 | 0.341 | 0.571 | 0.920 | 0.995 |
| 50 | | CompDTU | 1.00 | 0.049 | 0.062 | 0.115 | 0.215 | 0.370 | 0.720 | 0.931 |
| 50 | 50 | CompDTUme | 1.00 | 0.052 | 0.067 | 0.168 | 0.340 | 0.568 | 0.917 | 0.994 |
| 50 | 100 | CompDTUme | 1.00 | 0.051 | 0.067 | 0.166 | 0.340 | 0.569 | 0.917 | 0.995 |
| 50 | | CompDTU | 2.00 | 0.047 | 0.057 | 0.089 | 0.167 | 0.275 | 0.577 | 0.831 |
| 50 | 50 | CompDTUme | 2.00 | 0.052 | 0.074 | 0.164 | 0.341 | 0.575 | 0.917 | 0.996 |
| 50 | 100 | CompDTUme | 2.00 | 0.050 | 0.072 | 0.161 | 0.337 | 0.574 | 0.918 | 0.996 |
| 50 | | CompDTU | 4.00 | 0.055 | 0.051 | 0.080 | 0.121 | 0.185 | 0.400 | 0.637 |
| 50 | 50 | CompDTUme | 4.00 | 0.054 | 0.073 | 0.176 | 0.346 | 0.578 | 0.913 | 0.994 |
| 50 | 100 | CompDTUme | 4.00 | 0.052 | 0.071 | 0.171 | 0.343 | 0.574 | 0.917 | 0.995 |
| 100 | | CompDTU | 0.00 | 0.052 | 0.084 | 0.313 | 0.655 | 0.895 | 0.998 | 1.000 |
| 100 | 50 | CompDTUme | 0.00 | 0.052 | 0.084 | 0.313 | 0.655 | 0.895 | 0.998 | 1.000 |
| 100 | 100 | CompDTUme | 0.00 | 0.052 | 0.084 | 0.313 | 0.655 | 0.895 | 0.998 | 1.000 |
| 100 | | CompDTU | 0.01 | 0.051 | 0.082 | 0.322 | 0.638 | 0.895 | 0.999 | 1.000 |
| 100 | 50 | CompDTUme | 0.01 | 0.051 | 0.084 | 0.324 | 0.644 | 0.897 | 0.999 | 1.000 |
| 100 | 100 | CompDTUme | 0.01 | 0.051 | 0.084 | 0.324 | 0.644 | 0.896 | 0.999 | 1.000 |
| 100 | | CompDTU | 0.50 | 0.049 | 0.076 | 0.242 | 0.515 | 0.786 | 0.990 | 1.000 |
| 100 | 50 | CompDTUme | 0.50 | 0.049 | 0.084 | 0.315 | 0.643 | 0.892 | 0.999 | 1.000 |
| 100 | 100 | CompDTUme | 0.50 | 0.049 | 0.084 | 0.316 | 0.644 | 0.891 | 0.999 | 1.000 |
| 100 | | CompDTU | 1.00 | 0.056 | 0.074 | 0.201 | 0.425 | 0.687 | 0.965 | 0.999 |
| 100 | 50 | CompDTUme | 1.00 | 0.054 | 0.088 | 0.321 | 0.647 | 0.894 | 0.998 | 1.000 |
| 100 | 100 | CompDTUme | 1.00 | 0.056 | 0.086 | 0.319 | 0.649 | 0.896 | 0.998 | 1.000 |
| 100 | | CompDTU | 2.00 | 0.048 | 0.069 | 0.158 | 0.316 | 0.537 | 0.886 | 0.990 |
| 100 | 50 | CompDTUme | 2.00 | 0.054 | 0.094 | 0.320 | 0.635 | 0.894 | 0.999 | 1.000 |
| 100 | 100 | CompDTUme | 2.00 | 0.052 | 0.094 | 0.315 | 0.634 | 0.895 | 0.999 | 1.000 |
| 100 | | CompDTU | 4.00 | 0.048 | 0.055 | 0.118 | 0.217 | 0.364 | 0.706 | 0.932 |
| 100 | 50 | CompDTUme | 4.00 | 0.056 | 0.094 | 0.312 | 0.639 | 0.895 | 0.998 | 1.000 |
| 100 | 100 | CompDTUme | 4.00 | 0.052 | 0.088 | 0.309 | 0.638 | 0.898 | 0.999 | 1.000 |

**Supplementary Table S4:** Power for the *Comp* and *CompDTUme* Methods with Increased Between Subject Variance. The effect size $E$ is given by the numerical column names and ranges from 1 to 2.

| $n$ | $M$ | Method | $B$ | 1 (Null) | 1.10 | 1.25 | 1.375 | 1.50 | 1.75 | 2.00 |
|---|---|---|---|---|---|---|---|---|---|---|
| 26 | | CompDTU | 1.00 | 0.050 | 0.057 | 0.085 | 0.122 | 0.185 | 0.398 | 0.643 |
| 26 | 50 | CompDTUme | 1.00 | 0.052 | 0.065 | 0.108 | 0.176 | 0.294 | 0.598 | 0.861 |
| 26 | 100 | CompDTUme | 1.00 | 0.052 | 0.064 | 0.106 | 0.175 | 0.289 | 0.599 | 0.861 |
| 26 | | CompDTU | 1.50 | 0.048 | 0.056 | 0.078 | 0.104 | 0.154 | 0.295 | 0.504 |
| 26 | 50 | CompDTUme | 1.50 | 0.052 | 0.057 | 0.091 | 0.129 | 0.204 | 0.408 | 0.663 |
| 26 | 100 | CompDTUme | 1.50 | 0.051 | 0.056 | 0.090 | 0.128 | 0.201 | 0.408 | 0.662 |
| 26 | | CompDTU | 2.00 | 0.052 | 0.053 | 0.071 | 0.088 | 0.127 | 0.244 | 0.407 |
| 26 | 50 | CompDTUme | 2.00 | 0.054 | 0.056 | 0.078 | 0.103 | 0.153 | 0.310 | 0.518 |
| 26 | 100 | CompDTUme | 2.00 | 0.054 | 0.055 | 0.076 | 0.102 | 0.153 | 0.310 | 0.517 |
| 26 | | CompDTU | 4.00 | 0.050 | 0.054 | 0.064 | 0.070 | 0.090 | 0.156 | 0.234 |
| 26 | 50 | CompDTUme | 4.00 | 0.052 | 0.054 | 0.063 | 0.072 | 0.097 | 0.171 | 0.267 |
| 26 | 100 | CompDTUme | 4.00 | 0.052 | 0.053 | 0.063 | 0.070 | 0.096 | 0.171 | 0.266 |
| 100 | | CompDTU | 1.00 | 0.056 | 0.074 | 0.201 | 0.425 | 0.687 | 0.965 | 0.999 |
| 100 | 50 | CompDTUme | 1.00 | 0.054 | 0.088 | 0.321 | 0.647 | 0.894 | 0.998 | 1.000 |
| 100 | 100 | CompDTUme | 1.00 | 0.056 | 0.086 | 0.319 | 0.649 | 0.896 | 0.998 | 1.000 |
| 100 | | CompDTU | 1.50 | 0.052 | 0.065 | 0.156 | 0.323 | 0.546 | 0.901 | 0.994 |
| 100 | 50 | CompDTUme | 1.50 | 0.051 | 0.074 | 0.208 | 0.444 | 0.708 | 0.976 | 1.000 |
| 100 | 100 | CompDTUme | 1.50 | 0.050 | 0.073 | 0.206 | 0.444 | 0.708 | 0.976 | 1.000 |
| 100 | | CompDTU | 2.00 | 0.052 | 0.062 | 0.132 | 0.261 | 0.448 | 0.814 | 0.974 |
| 100 | 50 | CompDTUme | 2.00 | 0.050 | 0.067 | 0.167 | 0.336 | 0.563 | 0.912 | 0.994 |
| 100 | 100 | CompDTUme | 2.00 | 0.051 | 0.066 | 0.169 | 0.335 | 0.563 | 0.913 | 0.994 |
| 100 | | CompDTU | 4.00 | 0.049 | 0.060 | 0.097 | 0.154 | 0.256 | 0.546 | 0.819 |
| 100 | 50 | CompDTUme | 4.00 | 0.048 | 0.060 | 0.104 | 0.172 | 0.291 | 0.610 | 0.879 |
| 100 | 100 | CompDTUme | 4.00 | 0.048 | 0.060 | 0.103 | 0.171 | 0.292 | 0.610 | 0.878 |

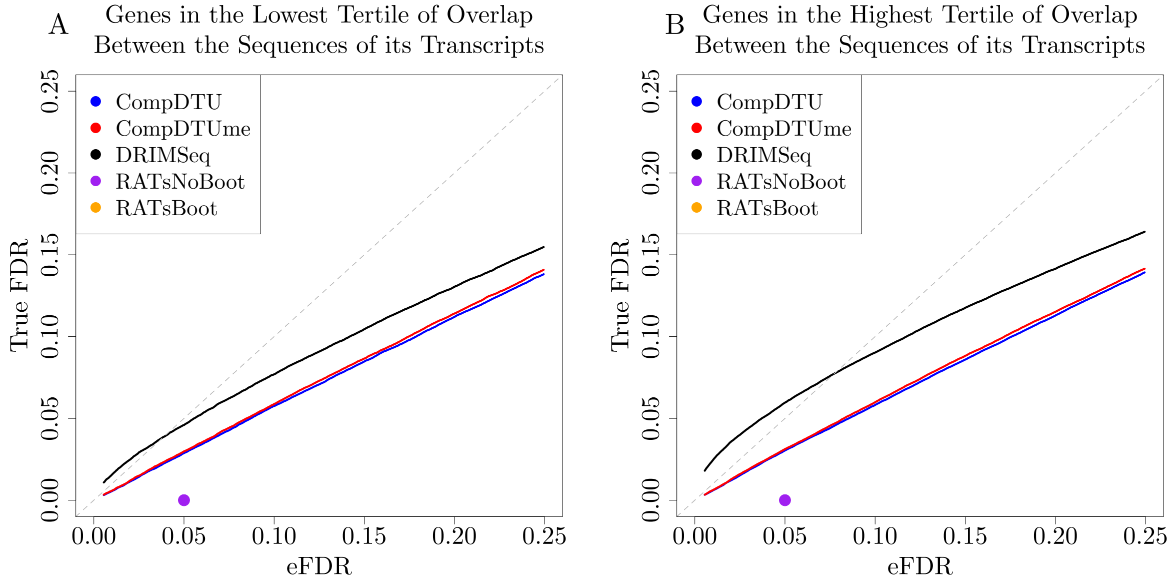

**Supplementary Figure S2:** Comparisons of false discovery rates under a moderate effect size (change value = 2). Both panels plot the True FDR level for a given eFDR value for the 7,522 genes that pass filtering in the analysis, where the eFDR is the FDR-adjusted $p$-value significance threshold. Panel A restricts to genes in the lowest tertile of sequence overlap, and Panel B restricts to genes in the highest tertile of sequence overlap. See Section 2 for details of how sequence overlap is calculated. *RATsBoot* is not shown in either panel because in Panel A the true FDR is 0.518 at an eFDR value of 0.05 and in Panel B the true FDR is 0.464 at an eFDR value of 0.05.

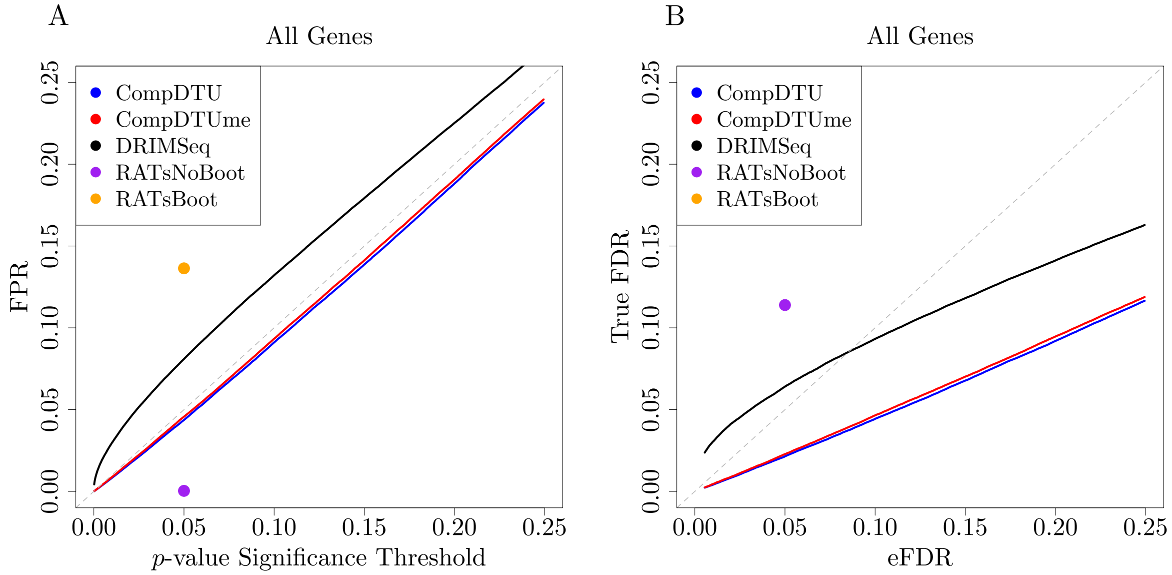

**Supplementary Figure S3:** Comparisons of false positive/false discovery rates under the null hypothesis (change value = 1, Panel A) and under a moderate effect size (change value = 2, Panel B). Panel A plots the $p$-value threshold used for rejection vs the false positive rate (FPR). Panel B plots the True FDR level for a given eFDR value, where the eFDR is the FDR-adjusted $p$-value significance threshold. Both panels plot the results for all 7,522 genes that pass filtering in the analysis.

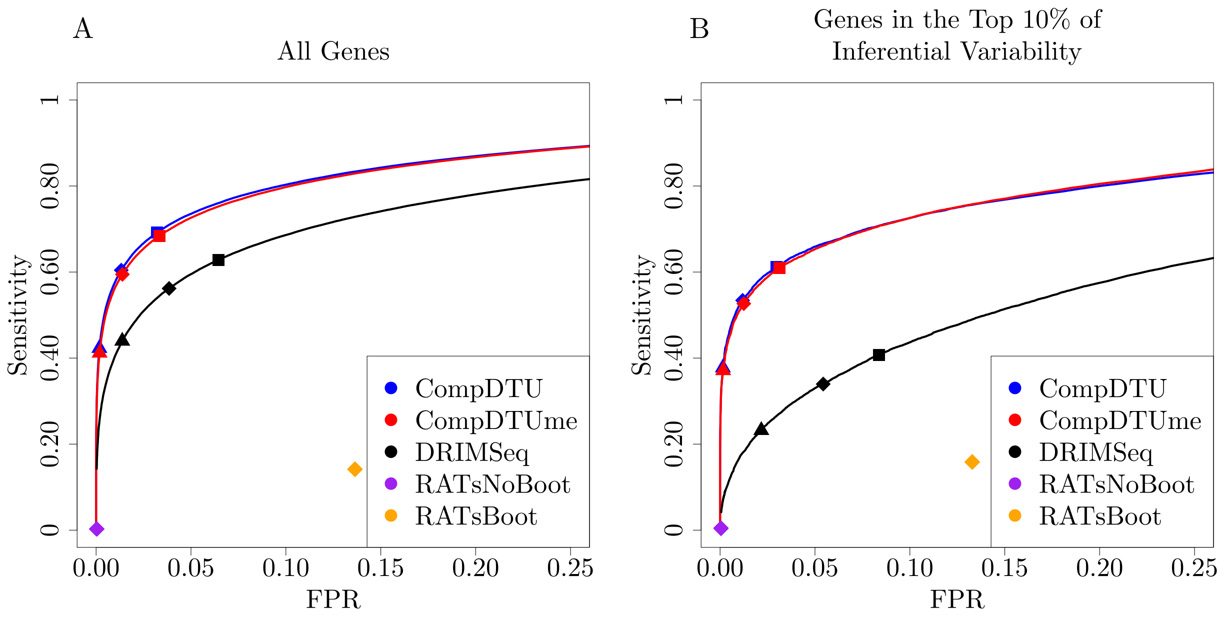

**Supplementary Figure S4:** ROC Curves for detecting DTU across 7,522 genes that pass filtering (Panel A) and the 752 genes in the top decile of gene-level inferential variability (Panel B). For details of how inferential variability is calculated, see Section 6 of the Supplementary Material (Van Buren and Rashid, 2020). A triangle, diamond, or square indicates the point at which the estimated FDR (eFDR) is 0.01, 0.05, or 0.10, respectively, where the eFDR is the FDR-adjusted $p$-value significance threshold.

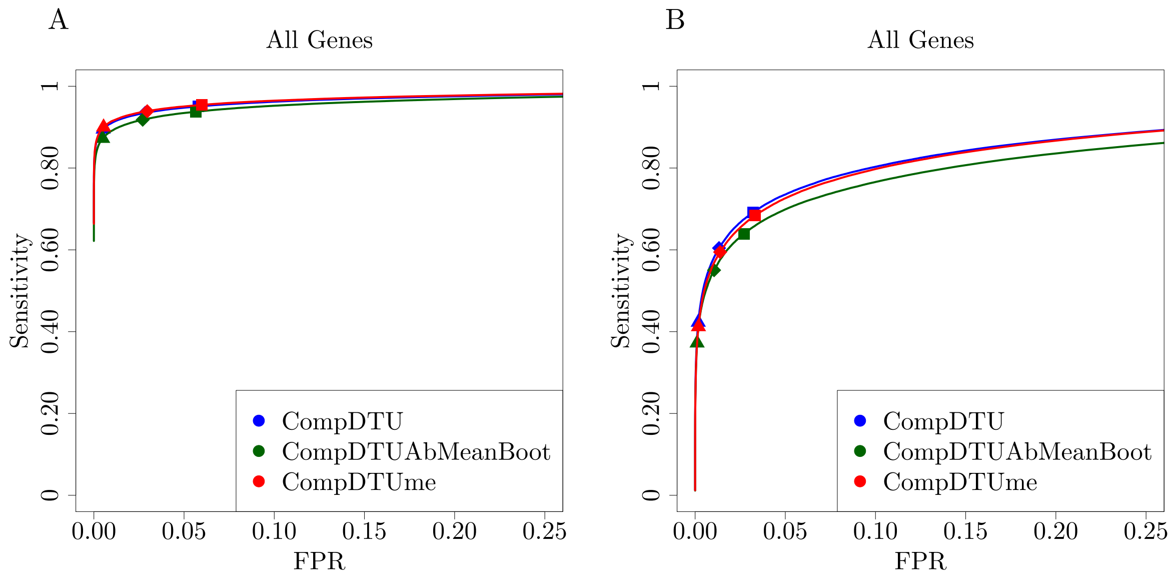

**Supplementary Figure S5:** ROC Curves for detecting DTU across the 7,522 genes that pass filtering for the 100 sample analysis (Panel A) and the 20 sample analysis (Panel B) including results for running *CompDTU* using the mean of the bootstrap samples instead of the usual point estimates (*CompDTUAbMeanBoot*).
